# Organ specific microenvironmental MR1 expression in cutaneous melanoma

**DOI:** 10.1101/2023.12.28.573554

**Authors:** Patricia B. Gordon, Woong Young So, Udochi F Azubuike, Bailey Johnson, James Cicala, Victoria Sturgess, Claudia Wong, Kevin Bishop, Erica Bresciani, Raman Sood, Sundar Ganesan, Kandice Tanner

## Abstract

The microenvironment is an important regulator of intertumoral trafficking and activity of immune cells. Understanding how the immune system can be tailored to maintain anti-tumor killing responses in metastatic disease remains an important goal. Thus, immune mediated eradication of metastasis requires the consideration of organ specific microenvironmental cues. Using a xenograft model of melanoma metastasis in adult zebrafish, we perturbed the dynamic balance between the infiltrating immune cells in the metastatic setting using a suite of different transgenic zebrafish. We employed intravital imaging coupled with metabolism imaging (FLIM) to visualize and map the organ specific metabolism with near simultaneity in multiple metastatic lesions. Of all the MHC complexes examined for brain and skeletal metastases, we determined that there is an organ specific expression of *mhc1uba* (human ortholog, *MR1*) for both the melanoma cells and the resident and infiltrating immune cells. Specifically, immune clusters did not express *mhc1uba* in brain metastatic lesions in immune competent fish. Finally, the differential immune response drove organ specific metabolism where tumor glycolysis was increased in brain metastases compared to skeletal and parental lines as measured using fluorescence lifetime imaging microscopy (FLIM). As MR1 belongs to the MHC class I molecules and is a target of immunotherapeutic drugs, we believe that our data presents an opportunity to understand the relationship between organ specific tumor metabolism and drug efficacy in the metastatic setting.

## Introduction

Cutaneous melanomas are characterized by elevated intratumoral, intertumoral and interpatient heterogeneities where a mosaic of cancer cells expressing mutated BRAF/ NRAS and/or the fully functioning form of the gene can be found within these lesions[1, 2]. Moreover, each clone may differentially express MHC I and/or MHC II proteins needed for antigen presentation for immune recognition and immune killing[3-5]. In some cases, multiple lesions can be detected within an organ concomitant with lesions in multiple organ sites such as the lung, liver, brain, intestines and bones[3, 4]. Inter-tumoral heterogeneity at each metastatic site further augments these genetic distinctions. This complexity is thought to blunt the efficacy of targeted therapies or combinatorial treatment with immunotherapies[3, 4]. Consequently, eradication of advanced melanoma in patients remains challenging [3, 4]. A major thrust in the field is to understand why therapies are ineffective, become ineffective, and show heterogeneous patient response in the metastatic setting.

The organ microenvironment regulates emergent tumoral genetic heterogeneities[6-8]. Far from being a static, steric hindrance, the dynamic reciprocal interactions provide environmental cues that bias survival of a subset of clones, evoke genetic changes of existing ones or is protective for some clones against therapeutic interventions[6-8]. Immune cells form an important component of the microenvironment and understanding how transformed cells continually escape immune targeted death remains a critical question[6, 7]. Moreover, these microenvironmental cues can drive the emergence of a broad range of functional states of infiltrating immune cells thereby complicating our ability to determine effective anti-tumor T cells[9-11]. This is especially pertinent to improve the efficacy of the class of immunotherapeutics in patient responders and non-responders[9, 10].

Here, we systematically examined the evolving heterogeneity of both the immune infiltrates and melanoma tumor clones in the presence and partial loss of T and B cells. Transcriptomic analysis was supplemented with direct visualization and metabolic mapping in the adult zebrafish model of melanoma metastasis to understand how tumors and their antigen presentation machinery are modified at different metastatic sites. Perturbation of the immune background of the transgenic fish resulted in a differential expression of cytotoxic T cells where CD8+ T cells were only found in the brain lesions in immune competent fish but not in skeletal lesions in either of the immune backgrounds. The model system also recreated the intratumoral genetic and MHC I/II heterogeneity observed in human patients which was validated using human orthologs of these genes from the TCGA database. We uncovered an organ specific expression of *MR1* (*mhc1uba*) where the relative expression of zebrafish gene *mhc1uba* was increased in the stromal cells as the immune infiltrates were decreased. This distinction in concert with the measured organ specific glycolysis illustrate that multiplexed methods can inform on microenvironmental cues that are important in our understanding of immune response.

## Results

The microenvironment is an important factor in defining intratumoral melanoma diversity[8, 12]. We examined the organ specific changes as a function of different adaptive immune microenvironments by comparing an immune competent zebrafish to the zebrafish equivalent of a SCID mouse: reduced T and B cells, [13]. Stromal cells such as neurons, fibroblast, leukocytes and keratinocytes were identified in both brain and skeletal lesions (Figure 1a). Reassuringly, glia cells were only found in the brain metastatic lesions independently of the immune background. Of all these components, leukocytes dominated the stromal components. We then stratified the different flavors of the immune infiltrates as a function of the immune microenvironment (Figure 1b). We identified cells that are constituents of innate and adaptive immune classifications. Focusing on the adaptive immune cells, increased numbers of cytotoxic (CD8+) T cells were identified in the brain lesions in immunocompetent fish compared to other conditions where very few cells were found in skeletal tissues (Figure 1b). Helper (CD4+) T and B cells were seen in comparable numbers conserved for each organ and immune background. However, on a per fish basis, helper T cells form the largest component of the immune infiltrates for all skeletal tumors. There was also an organ specific expression of Treg markers where the (CD8+) T cells found in the brain lesions did not express markers such as *CD25*, *CTLA4*, *IFN-γ* and *FOX3* (Supplemental figure 1). For the innate populations, two diverse sets of macrophages, *mpeg1.1*+ and *mfap4*+ were identified. *mfap4*+ cells were abundantly found in brain lesions in the immunocompetent fish compared to skeletal tumors in both backgrounds and to that of brain lesions in immunocompromised fish (Figure 1b). However, on average, similar numbers of *mpeg1.1*+ cells which clustered with dendritic cells were identified in all lesions (Figure 1b). Examination of the immune profiles demonstrated the variability in distributions of distinct immune populations on a per fish basis.

**Figure 1.**
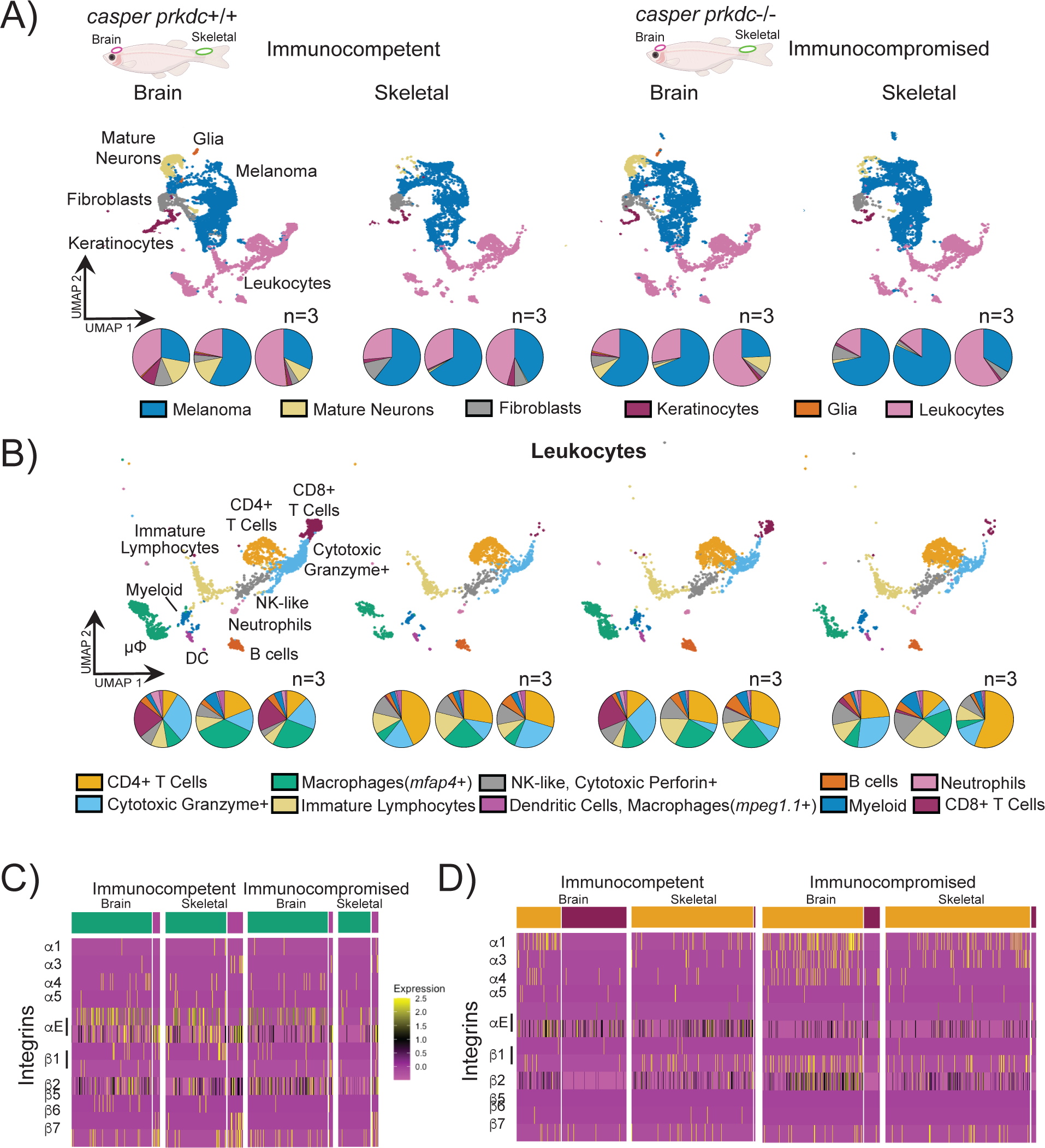
(A) UMAP of cells collected from melanoma tumors from brain and skeletal lesions from immune competent (*casper*) and immune compromised (*casper prkdc^D3612f^*) fish with distribution of cells, represented by the pie charts below of each individual sample. (B) The Leukocyte cells from the UMAP above showing the distribution of types of leukocytes with pie charts below showing the distribution of individual samples. (C) Heatmap of Macrophage clusters of integrin transcripts separated by immune background and organ. (D) Heatmap of T cell clusters of integrin transcripts separated by immune background and organ.

Immune infiltration requires that the cells possess molecular machinery underlying cell migration[6, 14, 15]. Moreover, integrin expression can further inform on anti-tumor effectiveness[16, 17]. Thus, we next quantified the expression of integrins and focal adhesion proteins of these infiltrating immune cells (Figure 1c and d, Supplemental Figures 1a, 2, and 3). Comparison of the different macrophage populations revealed that α3 integrin was highly expressed in the *mpeg1.1*+ macrophages/dendritic cells of skeletal tumors in each immune background. Cells within this cluster also co-expressed β6 and β7 integrins (Figure 1c, Supplemental figure 2). In contrast, *mfap4*+ macrophages showed the highest expressions of α4 (β2, β1) and β 7(αE2, αM) integrins in brain tumors (Figure 1c, Supplemental figure 2). In addition, β6 (α3) integrin was observed in *mfap4*+ macrophages for all tumors except for immunocompromised brain lesions (Figure 1c). Talin was the most abundant focal adhesion protein expressed for all sites of metastasis for both immune backgrounds (Supplemental figure 1a). A different integrin profile was quantified for the T cells. Alpha 1, 3 and beta 1 and 2 integrins and FAK were upregulated in CD4+ T cells compared to CD8+ T cells (Figure 1d, Supplemental figure 3). Talin was expressed in both types of T cells but in higher amounts for T cells found in immunocompromised brain and skeletal tumors (Supplemental figure 1a).

Melanomas are highly heterogeneous[4, 18]. Thus, we first asked if systemic immune modulation recreated this genetic diversity in metastatic lesions in this zebrafish xenograft model. Single cell transcriptomic analysis revealed seven distinct clusters in amelanotic brain and skeletal metastasis lesions (Figure 2a). Stratification based on previous studies revealed that clusters A, B, C, D, and G share Mature Markers, cluster E shares Stress Markers, and cluster F shares Neural Crest Markers (Supplemental figure 5)[19]. Examination of the distribution of each of these clones revealed organ specific distinctions. Cluster A was found in greater abundance in brain metastases compared to the skeletal metastatic lesions in the immunocompetent fish. In contrast, clusters E and F are found in greater abundance in skeletal lesions than those identified in the brain. Moreover, if we examine each fish, the relative abundance of each of these clones are maintained in each fish for most clusters with the exceptions of clusters C and D (Figure 2a).

**Figure 2.**
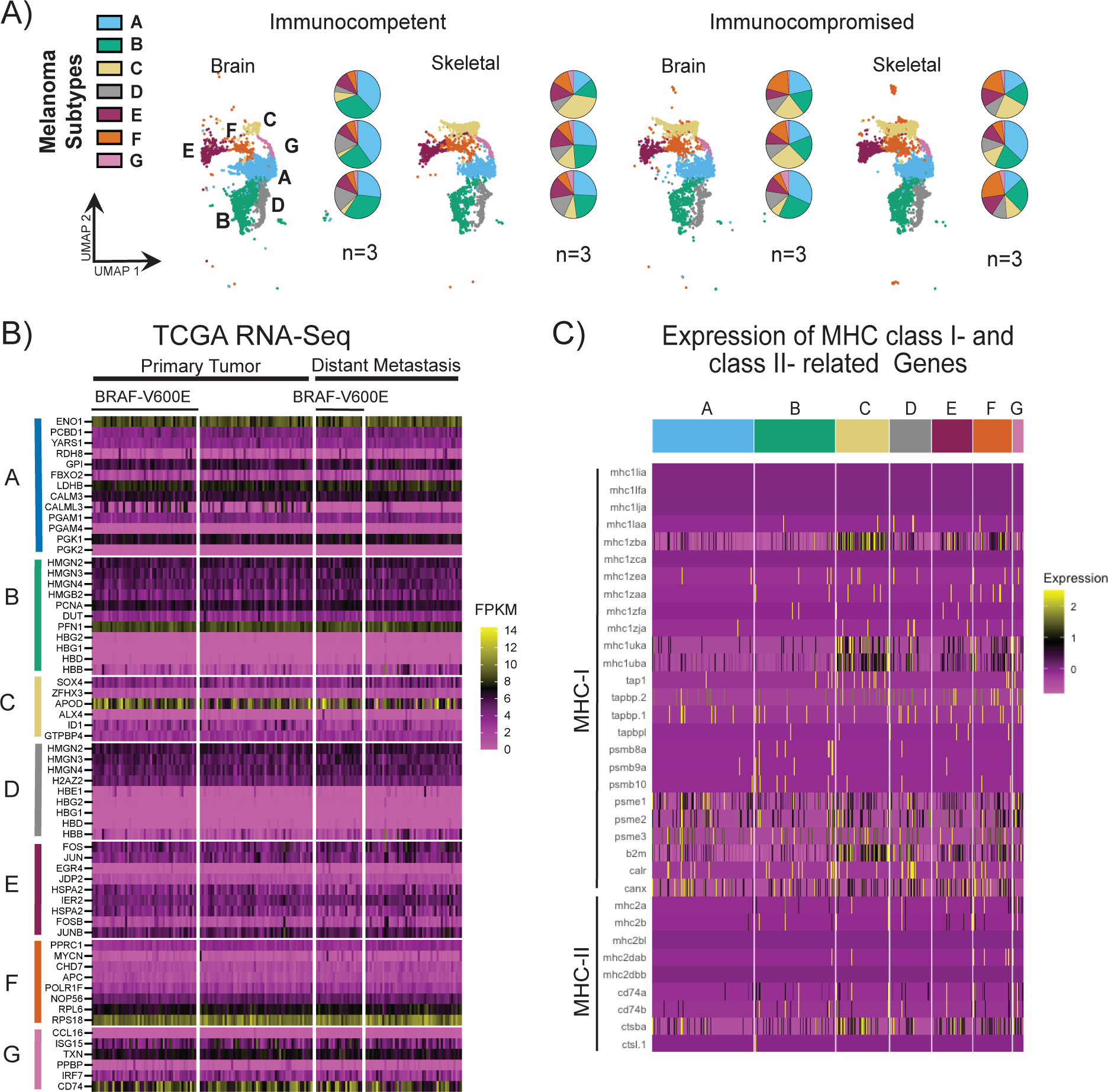
(A) The 7 melanoma clusters UMAP split by immune background and organ. Each UMAP represents 3 tumors and the pie charts below of the percentage of each cluster on a per fish basis. (B) A heatmap of the orthologues of the top 10 highest expressing genes from the zebrafish clusters using RNA-Seq analysis of TCGA-SKCM from samples of distant metastasis and primary tumors split by presence of BRAF-V600E mutational status. (C) Heatmap of MHC class I and class II expression from each melanoma cluster.

Infiltrating immune cells is known to sculpt the emergent genetic diversity, thus we next performed a direct comparison between immunocompromised and immune competent backgrounds[9]. We determined that the organ specific distribution of the clusters was skewed where the dominance of cluster A in the brain metastatic lesions was no longer observed. Instead, we observed increased representation of each cluster in each organ site. On a per fish basis, we observed variable distribution of melanoma clusters (Figure 2a).

To compare these clusters to genetic information obtained for human samples, we first siloed the publicly available data from the human skin cutaneous melanoma (SKCM) The Cancer Genome Atlas (TCGA) into mutational status (BRAFV600E vs. everything else) and primary vs. metastatic lesions for each of the ten top genes expressed in each of the aforementioned seven clusters (Figure 2b)[20]. We determined that these genes were expressed at values of >2 Fragments Per Kilobase of transcript per Million mapped reads (FPKM) for most of the top ten genes identified for each cluster. Focusing on the mostly highly expressed genes (>10 FPKM), we determined that genes identified in clusters A, B, C, F and G were highly expressed in the human data (Figure 2b). These genes are encoded for functions such as glycolysis (ENO1, cluster A), actin binding (PFN1, cluster B), stress in the aging human brain (APOD, cluster C), ribosomal (RPS18, cluster F) and antigen presentation (CD74, cluster G). Of all these genes, APOD and CD74 showed the greatest variation in expression for each tumor sample (Figure 2b). Thus, we next asked if the clusters showed distinctions in the antigen presentation machinery as a function of each cluster. We separated the genes associated with zebrafish antigens into two groups that are paralogs of the human major histocompatibility complexes I and II (MHC-I and MHC-II)[21]. Most genes are expressed at some level for each gene with notable exceptions. Some isoforms of MHC-I and MHC-II are not expressed in any cluster whereas the greatest expression of antigens was observed for clusters C and F while the converse was observed for cluster A (Figure 2c).

As CD8+ T cells target tumors by homing to peptide-MHC-I complexes derived via the MHC-I antigen presentation pathway, we next sought to understand what drove these distinctions in immune infiltration[22, 23]. As previously mentioned, zebrafish melanomas expressed many of the genes associated with antigen presentation (Figure 2c). We then asked if there was an organ specific distribution of MHC presentation for these amelanotic tumors. Of all the MHC proteins examined, transcriptomic analysis revealed an organ specific expression of *mhc1uba* (human ortholog, MR1) for the melanoma and immune clusters (Figure 3a, Supplemental figure 7). Moreover, immune clusters did not express *mhc1uba* in brain metastatic lesions in immune competent fish.

**Figure 3.**
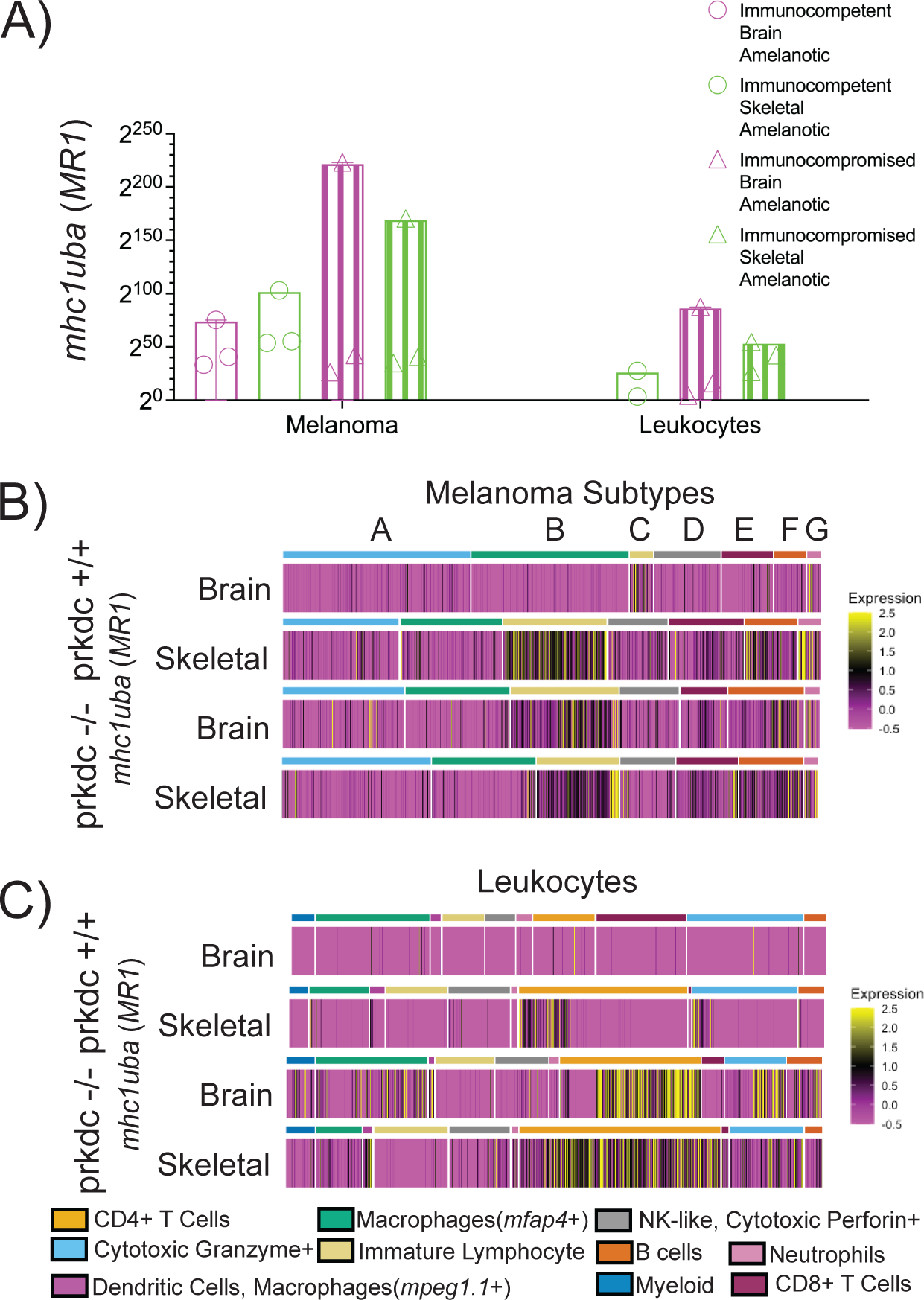
(A) Expression of *mhc1uba* (human ortholog *MR1*) in melanoma clusters and leukocyte clusters as a function of immune background and organ. (B) Heatmap of expression of *mhc1uba* in each cell organized by melanoma subtype for different immune backgrounds and organ. (C) Heatmap of expression of *mhc1uba* in each cell organized by type of leukocyte for different immune background and organ.

We then asked if there was cluster specificity for expression of *mhc1uba* (human ortholog, MR1). MR1 expression in clusters C, F and G were the major drivers of the increased expression of MR1 in skeletal tumors compared to brain tumors in immune competent fish (Figure 3b). These clusters also underpinned the overall increase for the immune compromised fish as these clusters were found at increased percentages for both the brain and skeletal tumors (Figure 3c).

In immune competent skeletal tumors, a small fraction of the CD4+ T cells and cytotoxic granzyme+ leukocytes were positive for MR1 expression (Figure 3c). Additional analysis confirmed that MR1 was not expressed by any immune or stromal cell populations in brain lesions in immunocompetent fish (Figure 3a and c, Supplemental figure 7). Instead, increased expression in CD4+ T cells, cytotoxic granzyme+ leukocytes in addition to other flavors of immune cells such as both *mpeg1.1*+ and *mfap4*+ positive macrophages and dendritic cells and B cells in immunocompromised lesions underpinned the overall increased expression observed for the immune compromised tumors seen in figure 3a (Figure 3c).

Macrophages have been shown to shape the metabolic landscape in metastatic melanoma lesions[24]. As we observed differences in MR1 expression in macrophages and other leukocytes, we next asked if there would be organ specific differences in metabolism as a function of the immune background. Fluorescence lifetime imaging microscopy (FLIM) can be used to directly image the abundance of the bound and free forms of reduced nicotinamide adenine dinucleotide (NADH) as each species has a different fluorescence decay lifetime (∼ns) free (0.1-0.5ns) vs bound (1-5ns), respectively[25]. By quantitating the relative contributions of each of these states, FLIM allows the determination of the relative rates of glycolysis or oxidative metabolism (Figure 4a). For example, an increased fraction of a_1_ (free NADH) corresponds to increased glycolysis whereas the inverse corresponds to increased oxidative phosphorylation (Figure 4a). As a result, the average lifetime of NADH (*τ_m_* = *a*_1_*τ*_1_ + *a*_2_*τ*_2_) was longer for skeletal tumors compared to brain tumors for each immune background (Figure 4b, Supplemental figure 8a and b). This can then be correlated to a metabolic state where FLIM analysis determined that brain melanomas had increased glycolysis compared to skeletal melanomas (Figure 4c and d). We then asked if FLIM could resolve metabolic differences linked to intratumoral heterogeneity within the organ. Comparison between brain lesions for each immune background revealed spatial heterogeneities in the measured lifetimes of both forms of NADH (Figure 4e and f). This was more pronounced for skeletal tumors where the spatial distribution of the lifetimes can be described by a bimodal gaussian distribution in the immune compromised background compared to the single gaussian distribution in the immune competent background.

**Figure 4.**
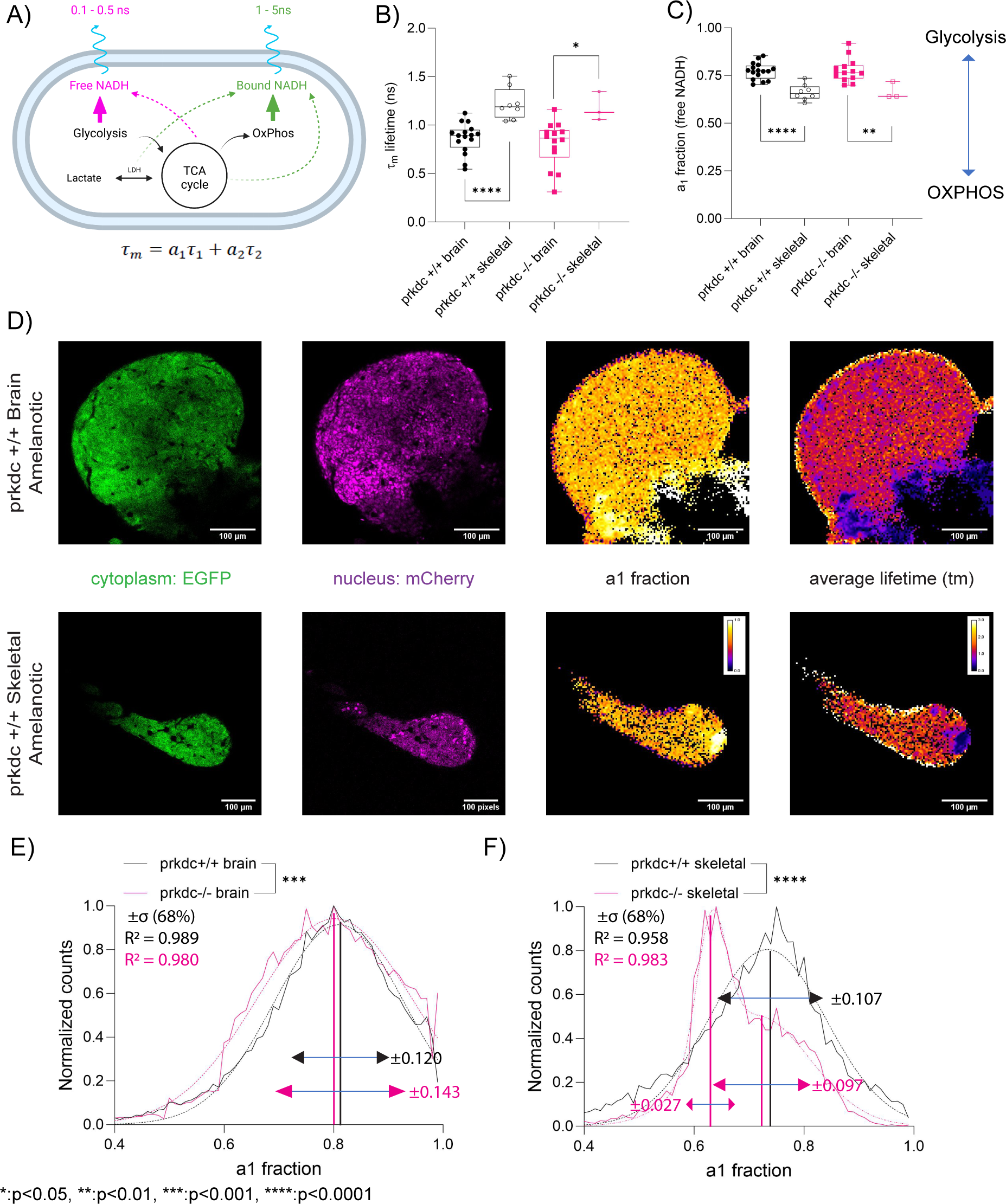
(A) Scheme of metabolic imaging to quantify the redox state of NADH to distinguish metabolism preference between glycolysis and oxidative phosphorylation (OxPhos) based on fluorescence lifetimes; free NADH as 0.1-0.5ns lifetime and bound NADH as 1-5ns. (B) Overall fluorescence lifetime (*τ*_m_) of NADH for melanoma tumor in terms of brain vs skeletal and *casper prkdc* +/+ vs *prkdc*-/-: *casper prkdc*+/+ brain (n=15), *casper prkdc*+/+ skeletal (n=8), *casper prkdc*-/- brain (n=13), and *casper prkdc*-/- skeletal (n=3). * p<0.05 and **** p<0.0001, unpaired two-tailed tests. Overall lifetime (*τ*_m_) was calculated based on free NADH fraction (a_1_) x lifetime of free NADH (*τ*_1_) + bound NADH fraction (a_2_) x lifetime of bound NADH (*τ*_2_). (C) Free NADH fraction to determine the metabolism preference between glycolysis and OxPhos for melanoma tumor in terms of brain and skeletal and *casper prkdc*+/+ and *casper prkdc*-/-: *casper prkdc*+/+ brain (n=15), *casper prkdc*+/+ skeletal (n=8), *casper prkdc*-/- brain (n=13), and *casper prkdc*-/- skeletal (n=3). ** p<0.01 and **** p<0.0001, unpaired two-tailed t-tests. (D) Images of *casper prkdc*+/+ amelanotic melanoma tumor in brain and skeletal, visualizing EGFP (cytoplasm), mCherry (nucleus), free NADH fraction (a_1_), and overall lifetime (*τ*_m_). (E) Overall histogram distribution of free NADH fraction (a_1_) between *casper prkdc*+/+ brain (n=15) and *casper prkdc*-/- brain (n=13) with single gaussian fit. *** p<0.001, two-way ANOVA. (F) Overall histogram distribution of free NADH fraction (a_1_) between *casper prkdc*+/+ skeletal (n=8) and *casper prkdc*-/- skeletal (n=3) with single gaussian fit for *prkdc*+/+ and bimodal gaussian fit for *prkdc*-/-. **** p<0.0001, two-way ANOVA.

## Discussion

Multiple factors contribute to the existence, emergence and persistence of genetic variances within a tumor[4]. A repertoire of driver mutations and environmental carcinogenic insults underpin multiple tumor clones in melanoma[4]. Additionally, microenvironmental factors can dominate cell fate decisions such as proliferation, metabolism and migration[1, 9, 26]. They can be siloed into cellular and acellular cues. The former include the resident and infiltrating cells. The latter include the cytokines, extracellular matrices and proteases within the organ milieu. As genetic alterations define metabolic needs, microenvironmental cues will also be a potent regulator of tumoral diversity[1, 9, 26]. Consideration of the localized cues within each of these organs is needed to understand disease progression, drug efficacy and drug resistance for metastatic disease. Here, we employed simultaneous metabolic mapping and intravital imaging to understand the interplay between organ specific and systemic immune regulation and tumor heterogeneity.

Both innate and adaptive immunity are critical for effective eradication of tumors[27, 28]. The tumor microenvironment can be immunologically “hot”, “variable” or “cold” dependent on the presence or absence of infiltrating cytotoxic immune cells and the cytokine profile of the milieu[29, 30]. Of these categories, immunotherapies are thought to be most effective in hot tumors[30]. In spite of the observation that melanomas are typically hot tumors, only a subset of patients respond to immunotherapies[31, 32]. These findings underscore the need to identify additional biomarkers and co-factors that expand therapeutic success in more patients[23]. Recently, studies focused on additional immune components in the microenvironment have begun to address what may cause T cell dysfunction and loss of anti-tumor responses[31, 32]. Here, we observed that both the skeletal and brain lesions were “variable” immunologically in immune competent fish. Notably, only cytotoxic T cells were found in the brain lesions. These cells may be exhausted or due to presence of additional cells such as myeloid cells may contribute to an immune suppressive environment in the brain. Moreover, comparison to tumors in the “SCID” background not only increased the diversity of the immune infiltrates, but immune cells also displayed different organ specific integrin profiles. These results provide a systematic way to interrogate immune modulation as a novel class of immunotherapies directly or indirectly targeting integrins are currently in varying stages of clinical trials for melanoma[33, 34]. Specifically, a small molecule named 7HP349 showed allosteric activation of α4β1 and αLβ2 integrins in T cells to increase T cell infiltration into colorectal and melanoma tumors and has entered into phase I trials [34, 35]. In our work, we observed differences in the expression of these integrins in T cells and macrophages, but additional work will be needed to identify the preferred, functional heterodimers. Nevertheless, targeting α4β1 may also have off target effects depending on the site of the lesion due to expression on other types of infiltrating or resident cells.

Microenvironmental factors convey epigenetic and transcriptomic alterations to drive increased intratumoral heterogeneities in melanomas[36]. The presence of these multiple tumoral states has been implicated in therapeutic resistance in primary and metastatic disease[36]. Moreover, the spatio-temporal changes may be regulated by organ specific cues from the tumor microenvironment[6, 37]. Thus, the subclonal evolution of cancer cells within the brain are driven by different environmental stressors from that within the bone further complicating treatment in patients with multiple lesions. Single cell transcriptomics has allowed for dissection of the intratumoral heterogeneity[38]. Multiple cell states in melanoma have been identified including neural crest, pigmented, mature, stress, invasive, and starved states[19, 38]. Here, our organ specific comparison also identified melanoma clones that exhibit transcriptomic markers associated with these classifications albeit with different organ specific ratios. Perturbation of the immune background increased the diversity of the clusters increasing the heterogeneity of the number and relative abundance of the clusters. These changes demonstrate the importance of both the immune environment and organ specific cues in driving intratumoral heterogeneity.

MHC class I and class II genes encode proteins that present antigens to T cells to aid in recognition of bacteria, viruses, and cancers[39]. Tumor cells can successfully evade immune surveillance by downregulation of antigen presentation machinery via modulating the expression of MHC-I and/or -II proteins[39, 40]. Thus, examination of the MHC-I and -II proteins can in part inform immunotherapy response[39, 41]. There is great conservation of these complexes between zebrafish and humans[21]. Moreover, the zebrafish tumors show heterogenous expression of MHCs comparable to human tumors where we observed an organ specific expression of *mhc1uba* (human ortholog, MR1) for the melanoma and immune clusters. None of the immune cells expressed MR1 in brain metastatic lesions in immune competent fish. MR1 has been implicated in riboflavin derived antigens from bacteria to Mucosal-associated Invariant T (MAIT) cells[42]. MR1 presentation can also occur to MR1-restricted T (MR1T) cells[42, 43]. MR1 antigen presentation machinery involves vitamin B metabolites, namely derivatives of riboflavin and folate[43]. It has been postulated that some tumor-related antigens can bind to MR1 and active MAIT cell[42]. In the absence of microbial antigens, a MR1-T cell clone can target leukemia and melanoma cell lines via tumor secreted MR1 molecules[44]. The cells can easily be detected in the blood of healthy individuals and were classified as a new cell population based on their capacity to recognize MR1 and on their ability to react to different types of cancer cells.

The dynamics of the immune microenvironment span different spatio-temporal scales co-operatively dictated by local resident cells and by the infiltrating cells[45]. As an example, within the brain microenvironment, resident macrophages such as microglia are restricted to immune surveillance within the brain[46]. However, infiltrating innate and adaptive immune cells are recruited from extracranial tissues at different times during tumor initiation and progression[47, 48]. These interactions may be transient but can profoundly change the metabolic landscape of the tumor microenvironment. The relative size of zebrafish render it amenable to optical techniques to visualize single tumor cell and immune dynamics in multiple organs with near simultaneity[49-51]. FLIM has been used to probe macrophage metabolism in vivo during wound healing in the larval zebrafish [52]. Here, we employed this powerful technique to measure tumor metabolism of multiple lesions in juvenile fish. Metabolic rewiring in melanoma results in shifts in glycolysis and oxidative phosphorylation (OXPHOS). This in turn can drive an acidic microenvironment deprived of glucose with deleterious effects of immune infiltration and anti-tumor efficacy resulting in a low response rate to immunotherapy [24]. We observed that metastatic brain lesions showed higher glycolysis compared to skeletal lesions when probed directly *in vivo* for treatment naïve fish in both immune compromised and immune competent fish. In recent studies, increased OXPHOS was determined from comparing patient matched metastatic brain tumors and extracranial metastases following therapeutic interventions including immunotherapies. This was validated by digested tumor cells excised from human tumors intracranially and subcutaneously in immunocompromised mice [24]. Our observed differences can be due to the different methods used to analyze metabolomics where one relies on dissociated tissue and the second due to the different sites of organ comparisons. Moreover, FLIM allows for single cell resolution where we can quantitate the intratumoral heterogeneity. This further stratified the organ specific heterogeneities where two distinct metabolic populations were detected in skeletal tissues in immune compromised fish compared to the immune competent where the metabolic population with reduced glycolysis (increased OXPHOS) formed a larger fraction of the tumor than that of the population with higher glycolysis. Recently, in a mouse model of adoptive T cell therapies in GBM and melanomas, a drug that modulates metabolism of the introduced T cells improved tumor clearance and reduced adverse side effects[53]. As T cell activation and melanoma metastatic potential are in part regulated by metabolic factors, these findings highlight the importance of a multi-pronged approach to decipher the interconnected organ specific microenvironmental cues [26].

In conclusion, the zebrafish animal model was the first animal model that recapitulated human melanomas. In these studies, we add to the growing body of evidence for the relevance of this model system in understanding metastatic disease and provide multiplexed analysis to inform on immune regulation of metabolism and cell dynamics. These data are critical in understanding what is needed to understand what drives anti-tumor T cell responses and as a future goal to improve efficacy in a broader range of patients.

## Acknowledgments

This effort was supported by the Intramural Research Program of the National Institutes of Health, the National Cancer Institute. This work utilized the computational resources of the NIH HPC Biowulf cluster (http://hpc.nih.gov). Services were provided by the CCR Genomics Core at the National Cancer Institute and the CCR Single Cell Analysis Facility (SCAF) at the National Cancer Institute. All schemes were created with BioRender.com.

## Materials and Methods

### Zebrafish husbandry

Animal studies were conducted under protocols approved by the National Cancer Institute and the National Institutes of Health Animal Care and Use Committee. Zebrafish were maintained at 28.5°C on a 14-hour light/10-hour dark cycle. Larvae were obtained from natural spawning, raised at 28.5°C, and maintained in fish water (60mg Instant Ocean sea salt per liter of water). Larvae were checked regularly for normal development and water was changed daily. Regular feeding began at 5 dpf.

### Zebrafish lines

The *casper* zebrafish has been previously characterized [54]. The *prkdc*-null SCID zebrafish, *prkdc^D3612fs^*, has been previously characterized [55]. Both lines were a kind gift from David Langenau.

### Cell culture and allograft

ZMEL1 cells were a kind gift from Richard White. Culturing of ZMEL1 cells were previously described [56]. 95% confluent cells were detached using 10mM EDTA in PBS. Cells were washed with PBS and resuspended into a concentration of 1 million cells per 10 microliters and 1.9 nL was injected into the mid-brain parenchyma at 4 dpf under anesthesia (Tricaine, MS-222). Zebrafish were allowed to recover for 24 hours in fish water before being put on housing system and feedings began at 5 dpf. Fish were sacrificed at 3 to 6 weeks of age.

### Single cell RNA Sequencing

Fish at 4 weeks of age were sacrificed and tumors were excised. The excised tumors were dissociated using Bresciani, *et al.* protocol for dissociation with 40ul Collagenase 100 mg/ml per 460ul 0.25% trypsin-EDTA[57]. Dissociated cells were kept at room temperature and submitted to CCR Single Cell Analysis Facility (SCAF). At SCAF, the cells were spun down, washed once with PBS and then counted. The cells were loaded into the lanes according to the 10X Genomics 3’ Single Cell User Guide with one capture lane per sample targeting recovery of 6,000 cells per lane. Cell partitioning was completed with uniform emulsion consistency and reverse transcription PCR was run. All subsequent steps of library preparation and quality control were performed as described in the 10X Genomics 3’ Single Cell User Guide using v3.1. Sequencing was performed by the CCR Genomics Core. One Illumina NextSeq runs for GEX were performed per four samples. The standard 10X Genomics *cellranger* version 6.0.0 pipeline was used to extract Fastqs and 10X Genomics *cellranger* version 6.0.0 pipeline was used to perform data processing. Sequenced reads were aligned to a custom zebrafish reference sequence, GRCz11.103 genome build. Seurat version 4.3.0 [58] was used to analyze the output from 10X Genomics *cellranger* pipeline. The data was trimmed by removing cells with less than 200 genes detected and more than 2500 genes detected. Cells with more than 5% of the transcripts belonging to mitochondria genes were also trimmed. After all individual samples were trimmed, the data was integrated. After the Seurat sctransform normalization [59] was performed, integration features were selected. Linear Dimensional reduction was performed using principal components analysis and 40 dimensions was used to run non-linear dimensional reduction, Uniformed Manifold Approximation and Projection (UMAP). K-nearest neighbors (KNN) graph was constructed in the PCA space using 40 dimensions. Using the Louvain algorithm, clusters were identified with a resolution of 0.5. Clusters were systematically identified as different cell types using table 1.

**Table 1.**
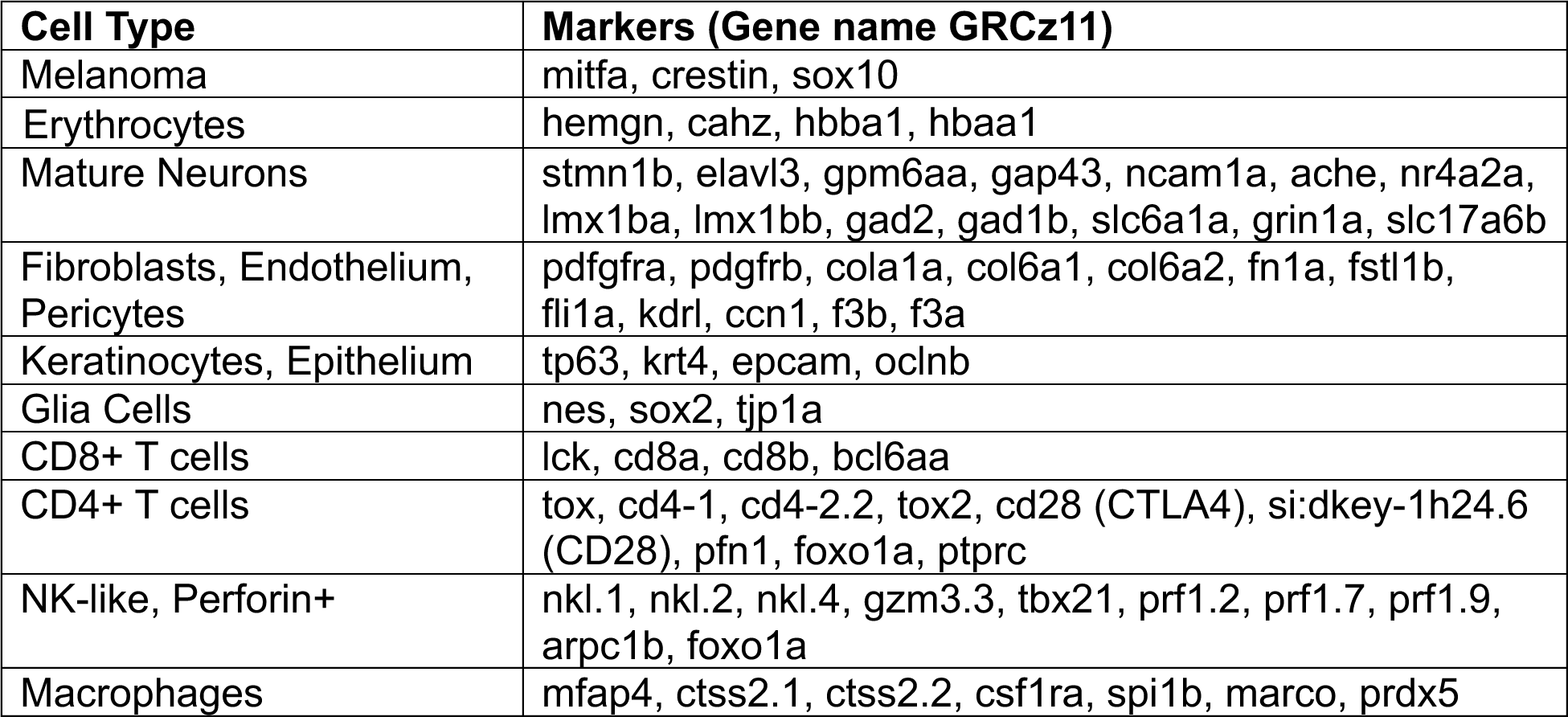
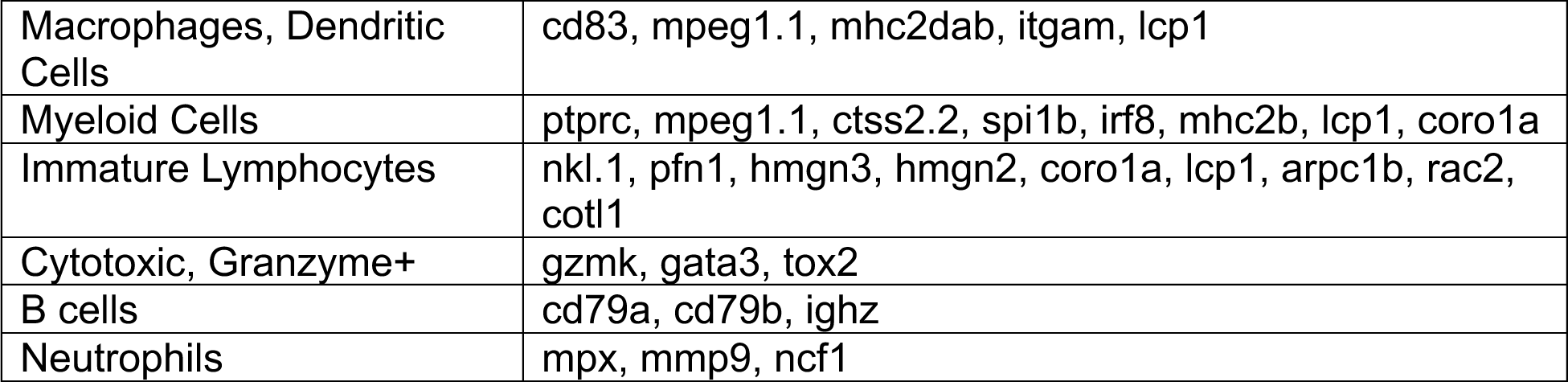

**Table 2.**
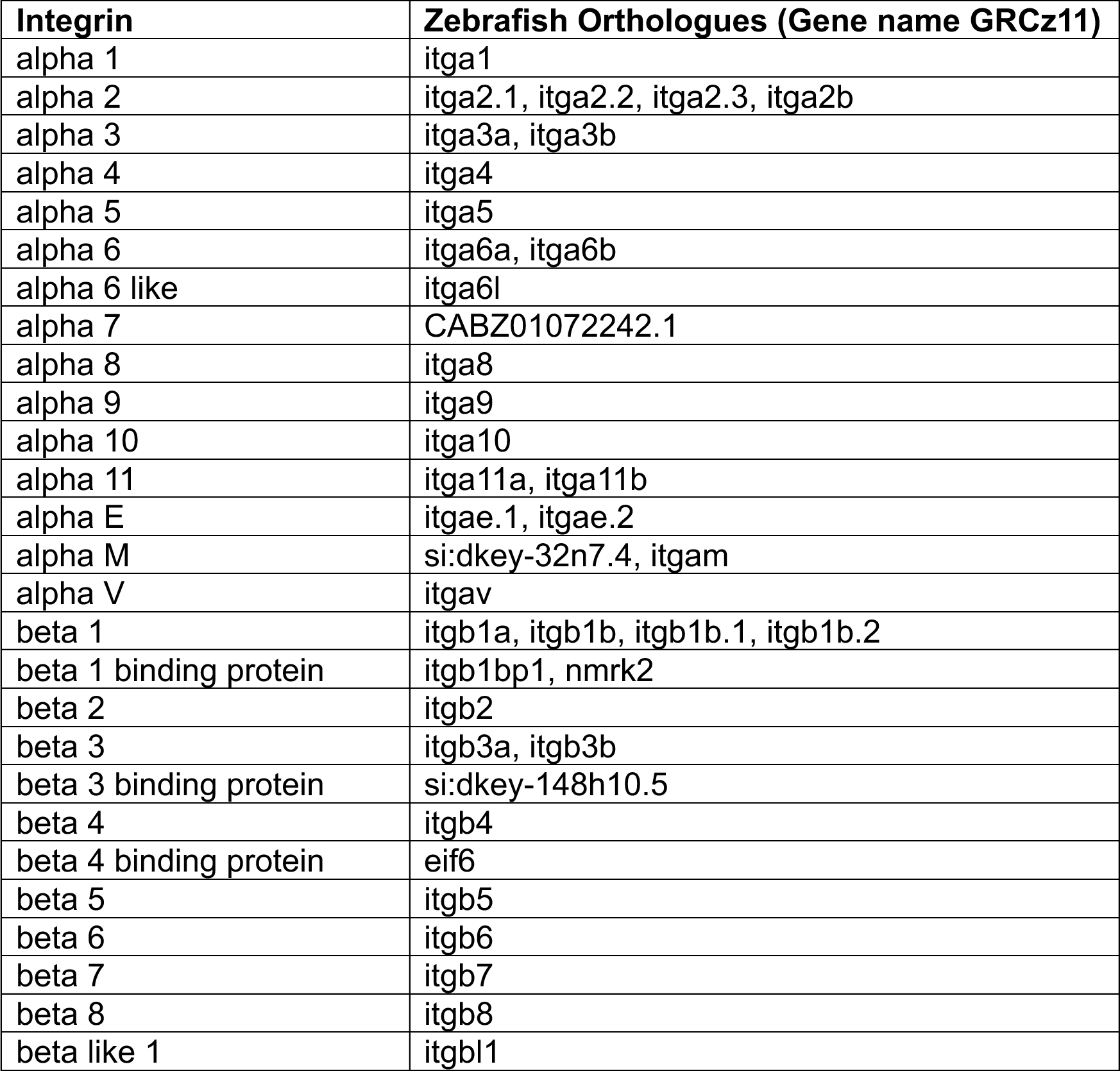

### Zebrafish and Human Homology

Genome build GRCh38 was used for all gene names for human genes and genome build GRCz11 for all gene names for zebrafish genes. Ensembl was utilized to find orthologues for human gene markers for specific cell types[60].

### Fluorescence lifetime imaging microscopy (FLIM) based metabolic imaging

FLIM was performed on melanoma tumors in the function of brain and tail *in vivo*. Fish with tumors were anesthetized with tricaine in system fish water (0.25x tricaine). Tricaine stock was prepared by dissolving 400 mg of Tricaine powder (ethyl 3-aminobenzoate methanesulfonate; Millipore Sigma, #E10521-50G) with 97.9 ml of deionized water and 2.1 ml of 1 M Tris. Anesthetic fish water was prepared by mixing 1.05 ml of tricaine stock per 100 ml of fish system water (0.1% buffered tricaine or 0.25x tricaine). Then, the tumors of anesthetized fish were imaged using a Leica SP8 WLL Falcon inverted confocal microscope with a 25x water immersive objective. A SP8 FLIM FALCON system was equipped with a tunable Multiphoton laser wavelength at 80MHz frequency and images were acquired as 512x512 pixel format with 1x zoom. As tumors had EGFP and mCherry, tumors were first excited at 800nm to capture both EGFP (510-550nm) and mCherry (580-640nm) simultaneously to correctly locate tumors of interest. After ensuring that fish did not move and were alive based on heart beating, the tunable multiphoton laser wavelength with 80MHz frequency was set to 750nm for metabolic imaging to capture the emission range of NADH (410-480nm) to delineate its state based on fluorescence lifetime decays in terms of free (0.1-0.5ns) and bound states (1-5ns). Fluorescence lifetime Images and lifetime decay traces were acquired and resolved by time-correlated single-photon counting using an SP8 FLIM FALCON system. Then, Images were fit, analyzed, and processed using LASX single molecule detection analysis software (Leica LASX) after a bin of 4 (2x2 surrounding pixels) to increase the fluorescence counts in each decay. A threshold or region was located based on EGFP and mCherry fluorescence of tumors. Then, fluorescence lifetime decays were fitted into a two-component exponential decay model,

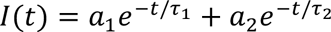

where a_1_ and a_2_ are the fractional contributions of the short and long lifetime components respectively (a_1_ + a_2_ = 1) and *τ*_1_and *τ*_2_ are the short and long lifetime components respectively. Short time components (a_1_ and *τ*_1_) corresponds to free NADH while long time components (a_2_ and *τ*_2_) are bound NADH.

---

**Supplemental Figure 1.**
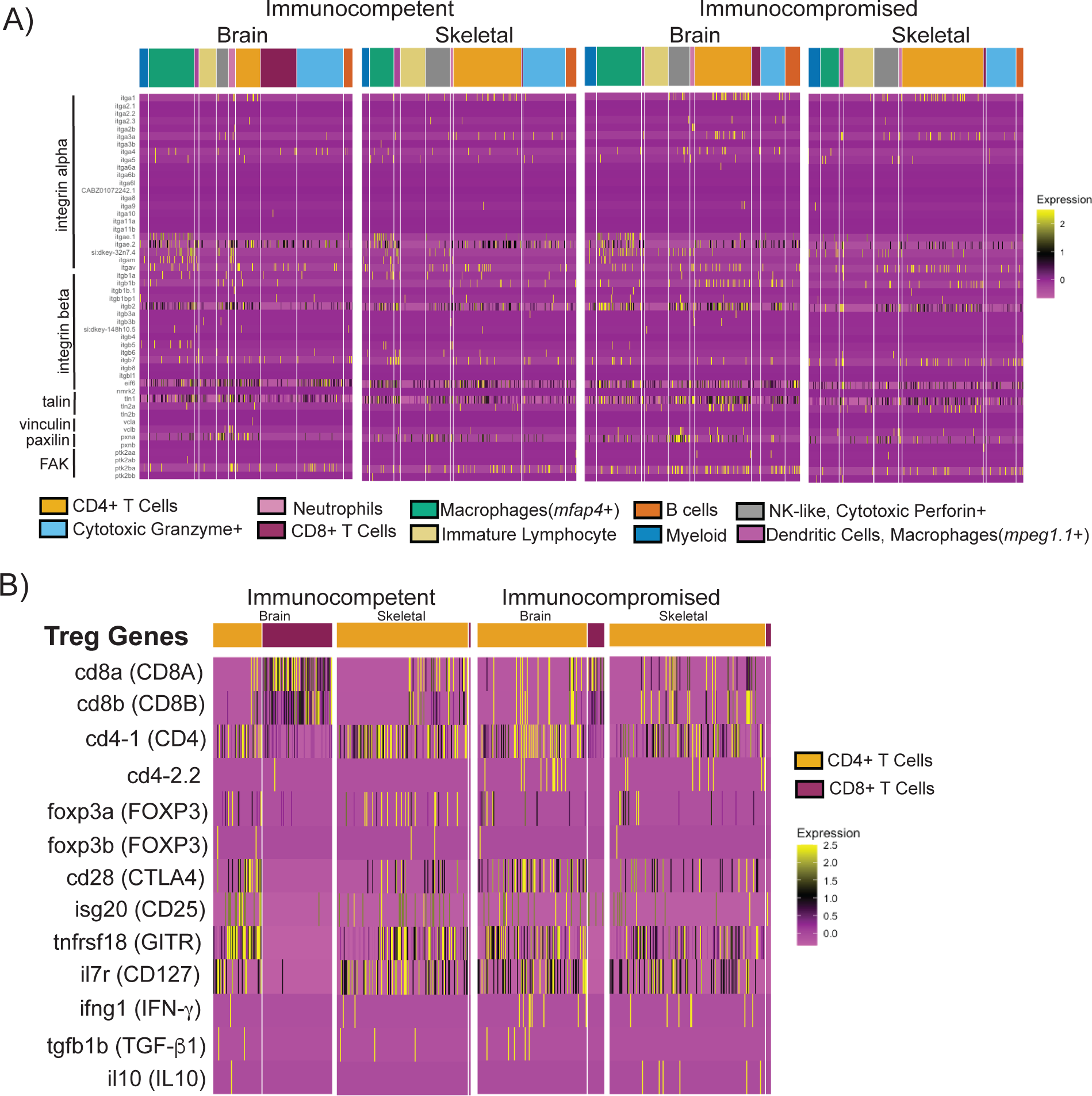
(A) Heatmap of alpha and beta integrins and focal adhesion transcripts in immune cell clusters, separated by immune background and organ. (B) Heatmap of several gene transcripts of genes associated with Treg cells of CD4+ and CD8+ T cells separated by immune background and organ.

**Supplemental Figure 2.**
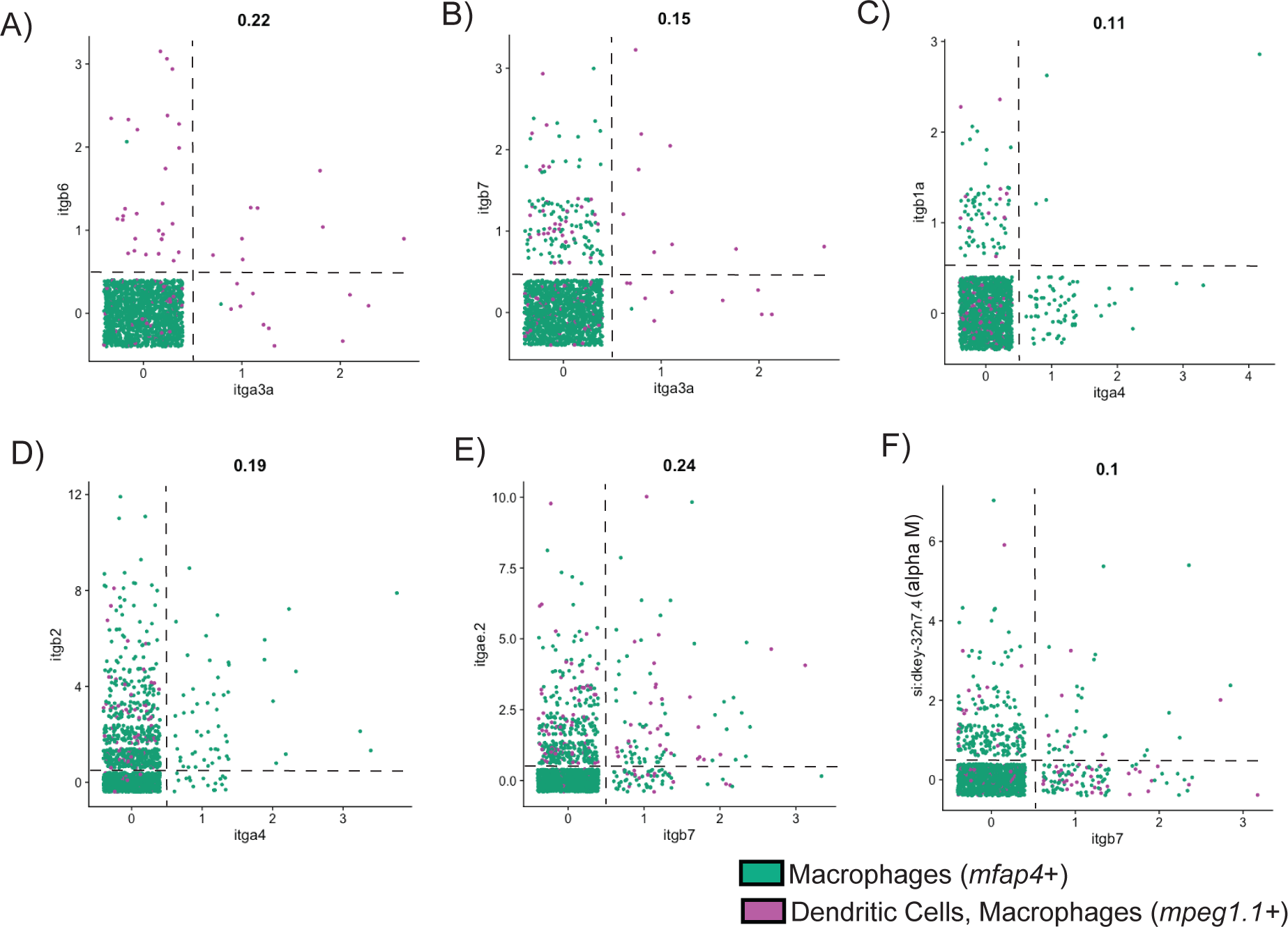
(A) Transcript correlation of integrins alpha 3 and beta 6 from macrophage cells (B) Transcript correlation of integrins alpha 3 and beta 7 from macrophage cells (C) Transcript correlation of integrins alpha 4 and beta 1 from macrophage cells (D) Transcript correlation of integrins alpha 4 and beta 2 from macrophage cells (E) Transcript correlation of integrins beta 7 and alpha E from macrophage cells (F) Transcript correlation of integrins beta 7 and alpha M from macrophage cells

**Supplemental Figure 3.**
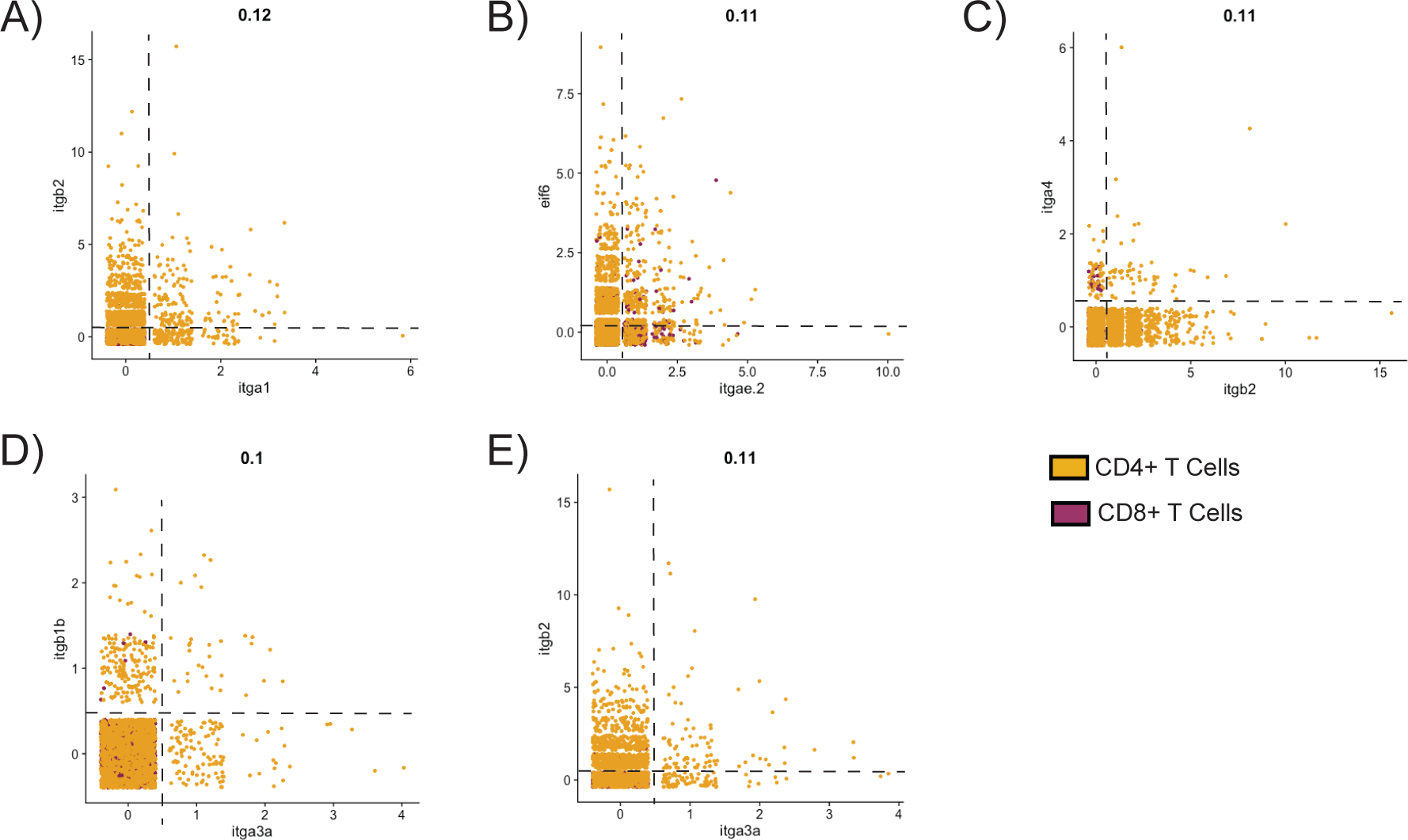
(A) Transcript correlation of integrins alpha 1 and beta 2 from T cells (B) Transcript correlation of integrins alpha E and beta 4 binding protein from T cells (C) Transcript correlation of integrins beta 2 and alpha 4 from T cells (D) Transcript correlation of integrins alpha 3 and beta 1 from T cells (E) Transcript correlation of integrins alpha 3 and beta 2 from T cells

**Supplemental Figure 4.**
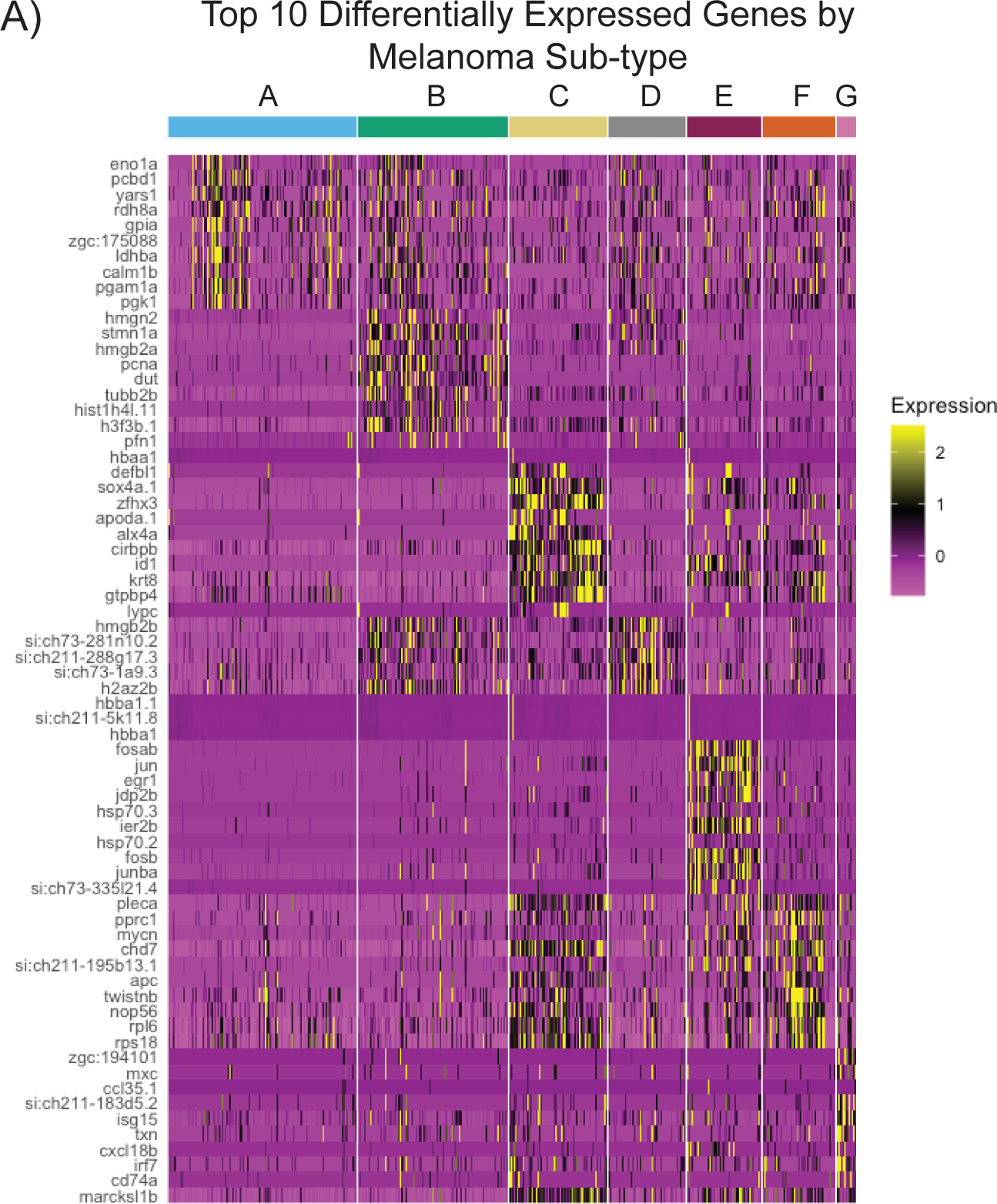
(A) Heatmap of the top 10 differentially expressed gene transcripts separated by cluster.

**Supplemental Figure 5.**
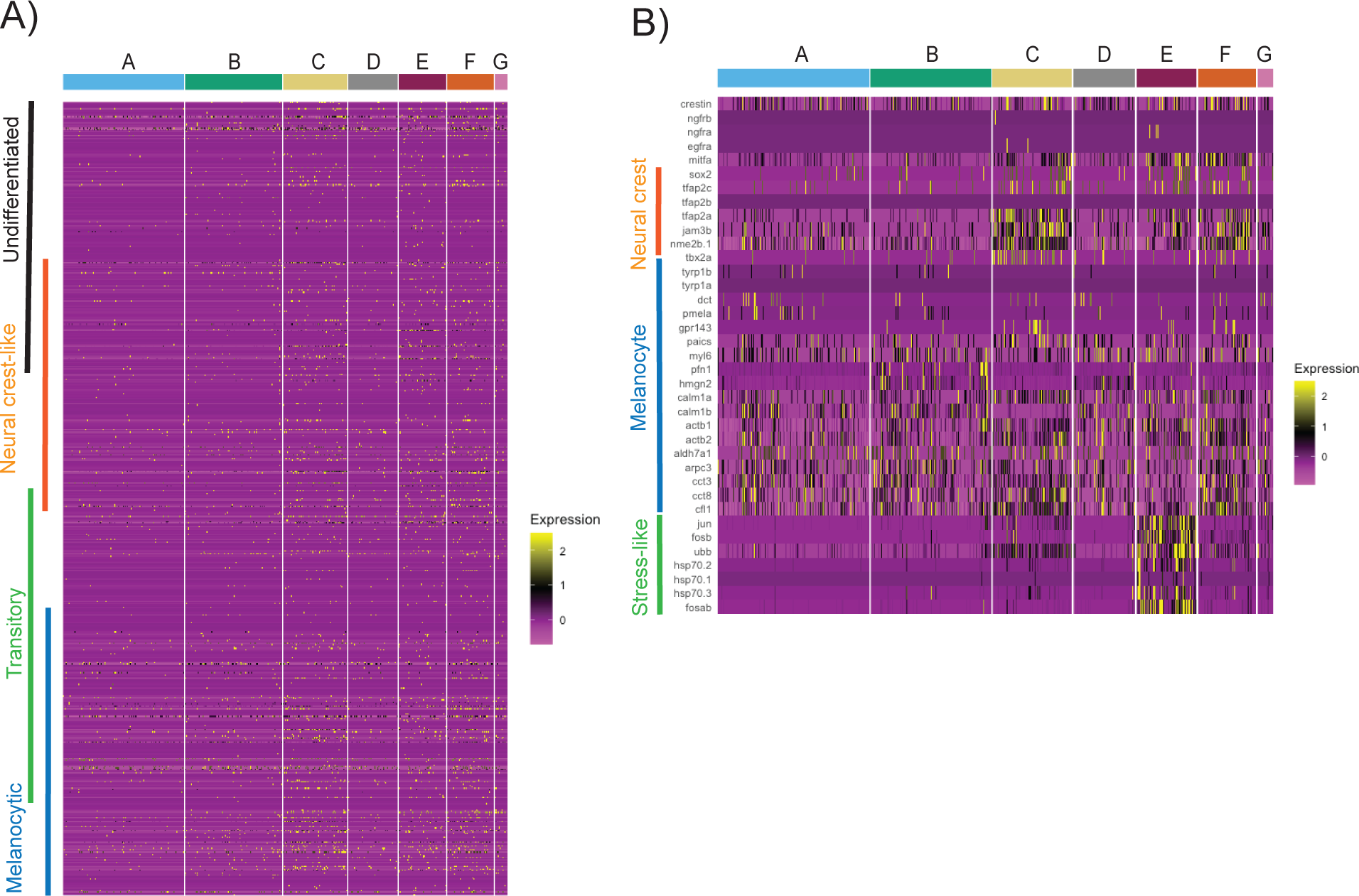
(A) Heatmap of gene expression from Tsoi, et al. 2018 melanoma subtype markers (B) Heatmap of gene expression from Baron, et al. 2020 of melanoma subtype markers

**Supplemental Figure 6.**
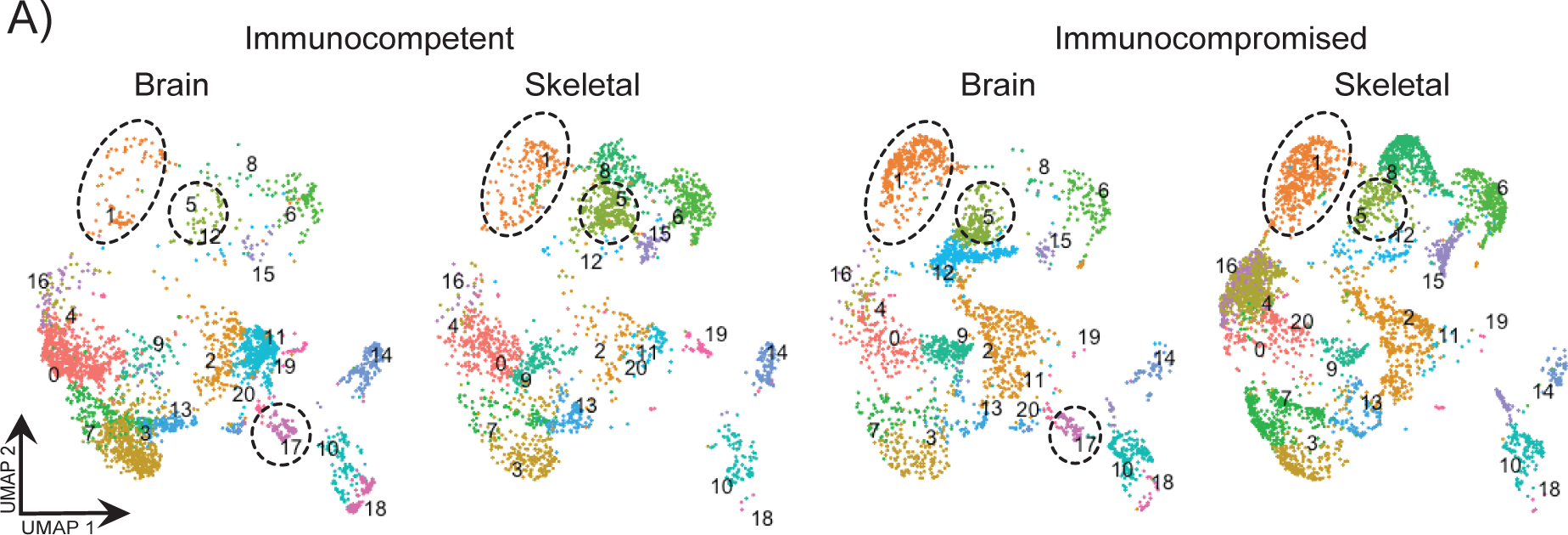
(A) UMAP of original melanoma clusters re-clustered, showing 21 different communities present, notably community 17 is not present in the skeletal samples and communities 1 and 5 are present at a diminished amount in the immunocompetent brain lesion

**Supplemental Figure 7.**
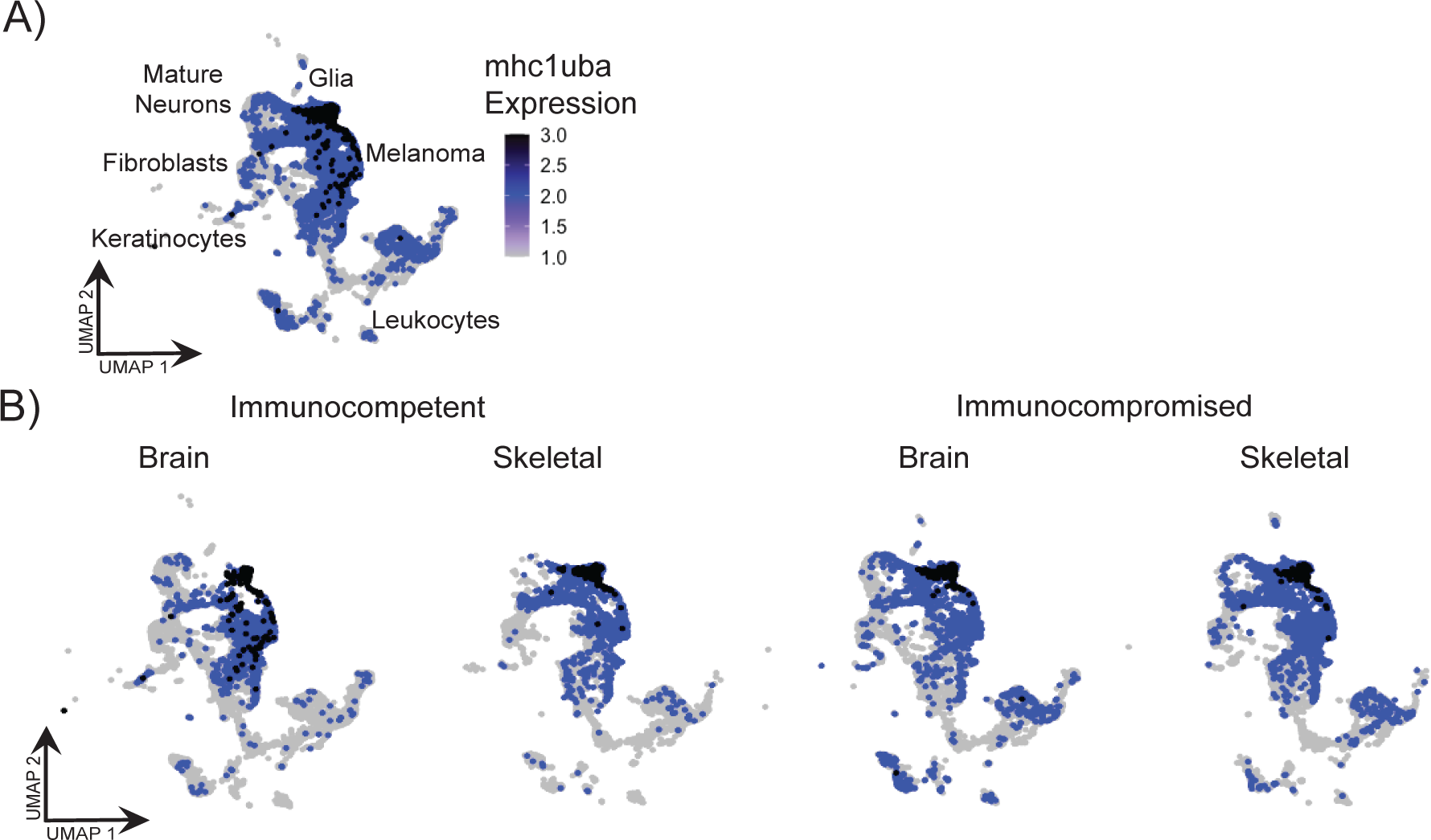
(A) UMAP of *mhc1uba* expression of *casper prkdc+/+* and *casper prkdc*-/- brain and skeletal melanoma samples (B) UMAP of *mhc1uba* expression of *casper prkdc+/+* and *casper prkdc*-/- brain and skeletal melanoma samples split.

**Supplemental Figure 8.**
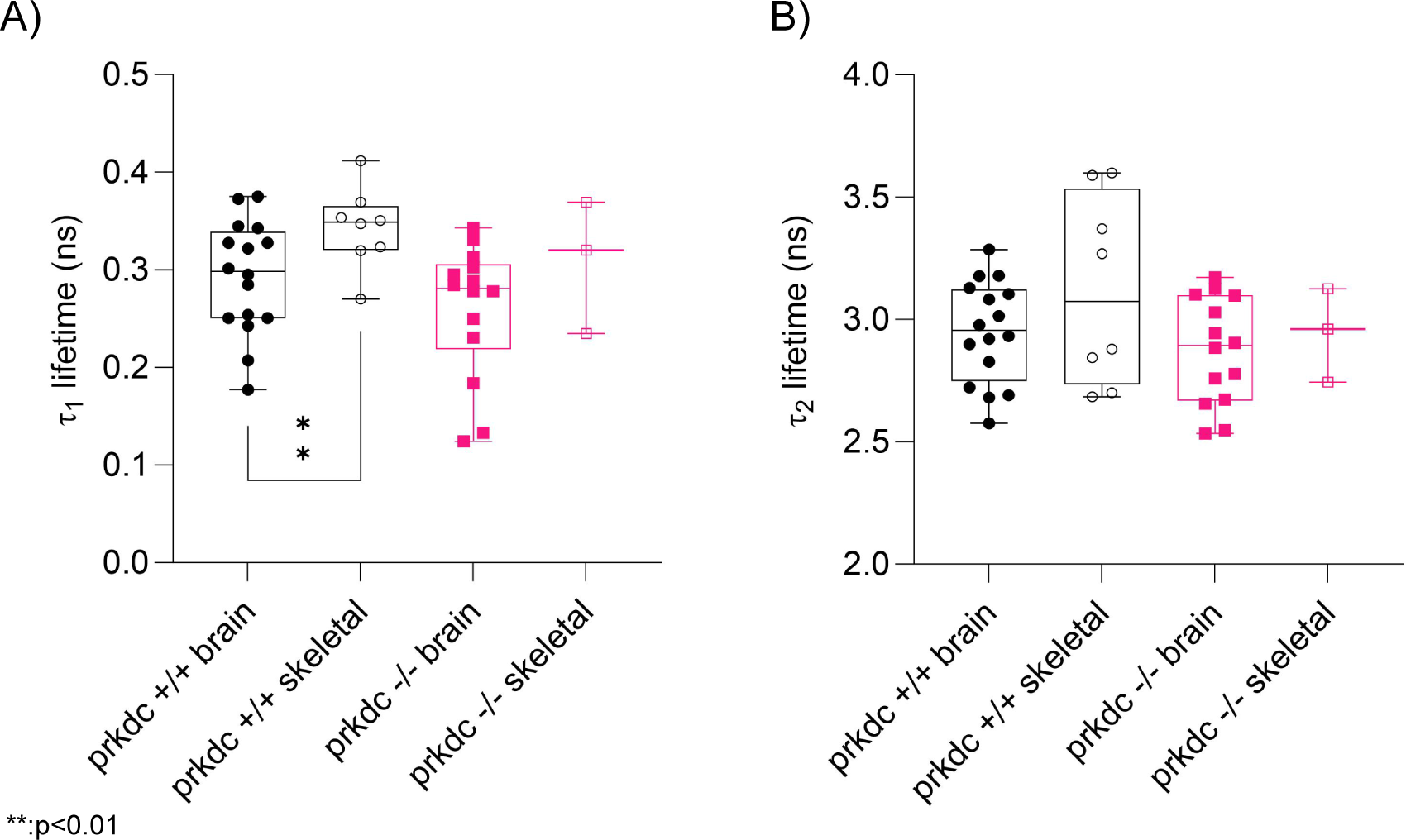
(A) lifetime of free NADH (*τ*_1_) for *casper prkdc*+/+ brain (n=15), *casper prkdc*+/+ skeletal (n=8), *casper prkdc*-/- brain (n=13), and *casper prkdc*-/- skeletal (n=3). ** p<0.01, unpaired two-tailed tests. (B) lifetime of bound NADH (*τ*_2_) for *casper prkdc*+/+ brain (n=15), *casper prkdc*+/+ skeletal (n=8), *casper prkdc*-/- brain (n=13), and *casper prkdc*-/- skeletal (n=3).

